# Spatial cueing effects do not always index attentional capture: Evidence for a Priority Accumulation Framework

**DOI:** 10.1101/2020.07.01.181826

**Authors:** Maya Darnell, Dominique Lamy

## Abstract

In visual search, improved performance when a target appears at a recently cued location is taken as strong evidence that attention was shifted to this cue. Here, we provide evidence challenging the canonical interpretation of spatial-cueing (or cue-validity) effects and supporting the Priority Accumulation Framework (PAF). According to PAF, attentional priority accumulates over time at each location until the search context triggers selection of the highest-priority location. Spatial-cueing effects reflect how long it takes to resolve the competition and can thus be observed even when attention was never shifted to the cue. Here, we used a spatial-cueing paradigm with abruptly onset cues and search displays varying in target-distractor similarity. We show search performance on valid-cue trials deteriorated the more difficult the search, a finding that is incompatible with the standard interpretation of spatial-cueing effects. By using brief displays (Experiment 1) and by examining the effect of search difficulty on the fastest trials (Experiment 2), we invalidate alternative accounts invoking post-perceptual verification processes (Experiment 1) or occasional failures of the onset cue to capture attention (Experiment 2). In Experiment 3, we used a combination of the spatial-cueing and dot-probe paradigms. We show that the events that occurred in both the cue and search displays affected attentional distribution, and that the relative attentional priority weight that accumulated at the target location determined how easily the competition was resolved. These findings fully support PAF’s predictions.

**Public significance statement:** Many studies aim at establishing whether certain objects mandatorily capture our attention. Here, we show that there is no “yes-or-no” answer to this question because the context in which an object appears determines whether this object captures attention. We show that our attention is not shifted to the highest-priority object at *any given time*: instead, information about priority is collected across time until some signal indicates that the appropriate moment for deploying our attention has arrived.

Striking failures to notice conspicuous events routinely illustrate how limited our attentional system is: we can attend to very few objects at any given time, and probably to just one. In natural conditions, when we move the focus of our attention from one object to another, we also shift our gaze towards the attended location: this allows us to place the object of most interest in the center of our fovea, which maximizes the quality of its perceptual processing. Tracking the locus of such *overt* attention is easily achieved by using eye-tracking devices. However, in order to isolate the benefits of attention from the benefits of visual acuity, one must study *covert* attention – that is, attentional shifts in the absence of eye movements. These shifts are not directly observable and must therefore be inferred using indirect measures of processing.

## The spatial cueing paradigm

A particularly popular method for studying covert attention is the spatial cueing paradigm. In a typical experiment, observers search for a target among distractors. Shortly prior to the search display, a cue appears at one of the potential target locations. Spatial cueing effects, that is, faster search performance when the target appears at the same location as the cue than at another location, are taken to indicate that attention was shifted to the cued location (see Eriksen & Hoffman, 1974; Posner, Nissen & Ogden, 1978; Posner, Snyder & Davidson, 1980 for early examples of this inference).

This method has been especially useful to investigate under what conditions an irrelevant object captures attention against the observer’s intention. For instance, Folk, Remington and Johnston (1992) asked participants to search for a color singleton target (e.g., a red target among gray distractors) in one condition, and for an abrupt onset, that is, a white target in an otherwise empty field, in a different condition. The location of the cue, also either a color singleton or an abrupt onset, was not predictive of the target location. Folk et al. (1992) observed a spatial cueing effect when the cue shared the target property (e.g., when it was an onset cue in search for an onset target), but not when it did not (e.g., when it was an onset cue in search for a color singleton target). This finding, which was replicated in numerous subsequent experiments (see Büsel, Voracek & Ansorge, 2018, for a meta-analytic review) led the authors to conclude that irrelevant salient objects do not capture attention unless they match the observers’ search goals.

Many authors have questioned the claim that failure to find a spatial cueing effect indicates that the cue did not capture attention. For instance, an alternative interpretation of Folk et al.’s (1992) finding is that a salient cue outside the observer’s attentional set captures attention, but attention is then quickly disengaged from its location during the cue-target interval. As a result, the spatial cueing benefit can no longer be observed when the target finally appears at the cued location (e.g., Theeuwes, Atchley & Kramer, 2000; but see Chen & Mordkoff, 2007; Lamy, 2005). Others have suggested that attention dwells at the location of the cue until the search display appears, and that the spatial cueing effect mainly indexes the time it takes to reject the distractor at the cued location (Gaspelin, Ruthruff & Lien, 2016). These authors thus proposed that irrelevant onset cues do capture attention, but their effects may or may not be observed: spatial cueing effects are reliable when the distractors are similar to the target and thus difficult to reject, but not when the distractors are dissimilar from it - as was typically the case in Folk and colleagues’ studies and their replications.

## Do spatial cueing effects index attentional shifts?

By contrast with the conflicting interpretations of null spatial cueing effects, the notion that, when found, spatial cueing effects indicate that attention was allocated to the cued location is fairly unanimous. However, we recently challenged this interpretation (Gabbay, Zivony & Lamy, 2019; Lamy, Darnell, Levi & Bublil, 2018). Relying on several observations, detailed below, we suggested that spatial cueing effects do not necessarily index attentional shifts.

Firstly, according to the standard explanation of spatial cueing effects, the cue is thought to capture attention before the search display appears. Thus, if the same cue produces no effect when target-distractor discriminability in the search display is high, but a large effect when such discriminability is low (Gaspelin et al., 2016; Lamy et al., 2018), one has to conclude that the cue captured attention in both cases. However, in order to explain why no spatial cueing is found in easy search (e.g., Folk et al., 1992), one must also assume that shifting attention takes no detectable time at all, even when the shift spans more than 10° of visual angle. This claim is difficult to reconcile with previous literature on attentional motion speed (e.g., Eriksen & Yeh, 1985; Tsal, 1983).

Secondly, spatial cueing effects have been reported with cue-target SOAs as long as 400ms (e.g., Gabbay et al., 2019). Although the time it takes to shift attention from one object to another during visual search is debated (e.g., Duncan, Ward, & Shapiro, 1994; Eimer & Grubert, 2014; Woodman & Luck, 1999), the most conservative estimates rarely exceed 200ms (e.g., Moore, Egeth, Berglan, & Luck, 1996). Finding a spatial cueing effect indicates that attention was not yet disengaged from the cued location when the search display appeared, according to the fast disengagement hypothesis (e.g., Theeuwes, 2010), or that it still dwelled at the cued location, according to the dwelling hypothesis (Gaspelin et al., 2016). Yet, it is difficult to explain why observers would not shift their attention away from a to-be-ignored cue for close to half a second, as both accounts would have to assume in order to explain the spatial cueing effects over long cue-target SOAs.

Thirdly, in a variant of the spatial cueing paradigm in which two successive cues rather than just one could appear prior to the search display, a performance benefit was observed at the locations of both the first and the second cue relative to uncued locations (Gabbay et al., 2019). Additional findings from Gabbay et al.’s (2019) study rule out the possibility that the residual benefit at the location of the first cue originated from trials in which the second cue failed to capture attention. This residual benefit is incompatible with the idea that attention was shifted from the first to the second cue and then to the target, as is typically assumed (e.g., Posner et al., 1980). If the spatial cueing effect is diagnostic of attention being allocated to the location of the cue, Gabbay et al.’s (2019) finding implies that attention was allocated simultaneously to two different loci (i.e., the locations of the first and second cues in the search display), a claim that is highly controversial (see Jans & Peters & de Weerd, 2010 for a review).

Finally, Lamy et al., (2018, Exp.1) recently conducted a variant of Gaspelin et al.’s experiment (2016, Exp.7), with three search difficulty levels: in the all-difficult condition, all distractors were similar to the target, in the all-easy condition they were all very dissimilar from it and in the mixed-difficulty condition, one distractor was similar to the target and two very dissimilar from it. We reported that, in line with the dwelling hypothesis (Gaspelin et al., 2016), an onset cue produced a spatial cueing effect that increased the more difficult the search. Critically however, we also found response times to increase with search difficulty when the target appeared at the cued location^1^, a finding that is incompatible with the canonic interpretation of spatial cueing effects. Indeed, this interpretation entails that faster responses at the cued location indicate that attention was shifted to that location and was still there when the target appeared, hence the benefit. Accordingly, responses to the target should be equally fast irrespective of what distractors (easy or difficult) surround the target.

## The Priority Accumulation Framework

In light of these observations, we proposed a priority accumulation framework (PAF, Gabbay et al., 2019; Lamy et al., 2018) that can accommodate the extant findings with no need to postulate cost-free shifts of attention, excessive attentional dwelling at irrelevant locations, or split foci of attention. By reinterpreting spatial cueing effects, which have long been the hallmark of attentional allocation, this account reconciles apparently conflicting findings, especially in the context of involuntary attentional capture.

According to PAF, the attentional priority accruing to a given location mainly depends on the physical salience of the successive objects that have appeared at that location (e.g., a cue, a distractor or the target) and on their similarity to the target, as well as on random noise. Priority weights accumulate over time at each location, until the search context signals that selection can occur. The first attentional shift is then made to the item that wins the competition (i.e., the item that has accumulated the highest attentional priority). However, how long it takes for the competition to be resolved varies as a function of how large the winner’s leading edge is. This is why on valid-cue trials, the competition is resolved faster the less similar the distractors are to the target.

A notable consequence of this scenario is that spatial cueing effects may occur even if attention was never directed to the cue: This may happen if the target is the highest-priority object both when it is cued and when it is not – for instance, when search is relatively easy and the cue’s weight is relatively small because it does not match the target-defining property. In that case, spatial cueing effects simply reflect that the competition that leads to target selection is resolved faster when the target location benefits from the extra-activation provided by the cue (valid-cue trials) than when it does not (invalid-cue trials). When search is very easy, competition can be resolved equally fast, irrespective of whether or not the target location is cued, hence the absence of any cueing effect. Finally, in difficult search, when the target is not the cued object, one or several distractors may successively become the focus of attention before the target is selected. In that case, the spatial cueing effect may reflect several additional processes including the time it takes to reject the distractor(s) and shift attention to the target.

We further suggested that unlike spatial cueing effects, effects of the compatibility between the response associated with the cued distractor and the response associated with the target attest that substantial perceptual processing took place at the cued location – provided that the discrimination that determines the appropriate response is difficult enough to require focused attention (e.g., Carmel & Lamy, 2014; Folk & Remington, 2006; Lamy et al., 2018). Thus, compatibility effects can provide reliable evidence that attention was allocated to the cued location^2^. In other words, a spatial cuing effect can be taken as evidence that attention was shifted to the location of the cue only if there is also an effect of the compatibility between the cued distractor and target response features. By contrast, a spatial cueing effect with no compatibility effect does not provide conclusive evidence for an attentional shift.

## Objective of the present study

Although PAF can explain most of the results that are problematic for the standard interpretation of spatial cueing effects, alternative accounts are possible. The objective of the present study was to assess these accounts and to provide new tests of PAF.

In Experiments 1 and 2, we focused on the finding that target-distractor similarity modulated performance on valid-cue trials (e.g., Lamy et al., 2018). Although as noted earlier, this finding appears to be at odds with the standard interpretation of spatial cueing effects, it is open to two alternative accounts that are compatible with that interpretation.

The first alternative account is that, as search displays were presented until the participants responded, the finding of a search-difficulty gradient on valid-cue trials may simply reflect processes unrelated to attention: when given enough time, participants may verify that the object at the cued location is indeed the target by comparing it to surrounding distractors, a process that should take more time when these are similar to the target than when they are not. Experiment 1 was designed to test whether we would replicate our findings when this verification process was prevented.

The second alternative account relies on the observation that the cue is likely not to capture attention on each trial. On trials in which the cue fails to capture attention, it obviously takes more time to find the target when search is difficult than when it is easy. This scenario entirely explains the search-difficulty gradient on valid-cue trials. In Experiment 2, we tested this account against PAF’s.

In Experiment 3, we sought more direct evidence for PAF’s claim that a cue can produce significant spatial cueing effects without ever becoming the focus of attention. We also tested PAF’s hypothesis that whether an object in the search display is the first to receive attention jointly depends on whether the cue appeared at its location, on the similarity of this object to the target and on how difficult the competition with other objects in the search display is to resolve.

## Experiment 1

Experiment 1 was similar to Lamy et al. (2018, Exp.1). Participants searched for a perfect circle among three elliptical shapes. A small black dot appeared on either the left or the right inside of each shape in the display. Participants had to report the side of the dot in the target circle. On any given trial, the search display included distractors that were all similar to the target (all-difficult search), all dissimilar from the target (all-easy search) or mixed (one similar and two dissimilar distractors - mixed-difficulty search). Prior to the search display, an abrupt onset was flashed randomly at one of the potential target locations. Unlike in previous versions of this task (Gaspelin et al., 2016; Lamy et al., 2018), the search display was presented briefly rather than until response. Responses were not speeded and accuracy was the dependent measure.

Our main interest was in how overall distractor-target similarity in the search display would modulate performance on valid-cue trials. We expected this modulation to disappear if it resulted only from post-attentional verification processes, and to remain significant otherwise.

We also expected to replicate the other findings reported by Lamy et al. (2018), which are compatible with both Gaspelin et al.’s (2016) dwelling hypothesis and PAF. Spatial cueing (henceforth, cue validity) effects should increase the more difficult the search. In the mixed-difficulty condition, accuracy should be lower when the difficult vs. an easy distractor is cued. Compatibility effects between the target and the cued distractor on invalid-cue trials should be observed only when a difficult distractor is cued. Finally, in the mixed-difficulty condition, we expected the response compatibility of the difficult distractor with the target to affect performance even when this distractor was not the cued object^3^.

## Methods

### Participants

We calculated the sample size required in order for the smallest meaningful effect reported by Lamy et al. (2018, Exp.1) to be significant, namely, the effect of the compatibility between the cued difficult distractor and the target in the mixed-difficulty condition. We conducted this analysis with G*Power (Faul, Erdfelder, Buchner & Lang, 2013), using an alpha of 0.01 and power of 0.95. We found the minimum sample size required to be 14 participants. We were therefore confident that our experiments would be sufficiently powered with a sample of twenty-one participants. All participants were undergraduate students (19 female, age: M = 23.34, SD =2.48) who participated for course credit. Participants reported normal or corrected-to-normal visual acuity. In this and the following experiment, all participants signed a consent form prior to the experiment. All protocols were approved by Tel Aviv University ethics committee.

### Apparatus

The experiment took place in a dimly lit room. Stimuli were presented on a 23-in. LED screen, using 1,920 × 1,280 resolution graphics mode and 120-Hz refresh rate. Responses were collected via the computer keyboard. Viewing distance was set at approximately 60 cm from the monitor using a chinrest.

### Stimuli

A sample sequence of events is presented in Figure 1. The fixation display consisted of five gray (138, 138, 138) square outline placeholders (2.4° x 2.4° of visual angle), one centered at fixation and the remaining four equally spaced at the corners of an imaginary square subtending 10° in diameter (i.e., central-frame center to outer-frame center distance was 5°). The onset-cue and search displays were similar to the fixation display, except for the following changes. In the onset-cue display, a cue consisting of four white dots (255, 255, 255; 0.5° in diameter) forming an imaginary diamond (3.3°x3.3°) was added around one of the four outer placeholders. In the search display, a filled red shape (255, 0, 0) appeared in the center of each of the outer placeholders: one circle (the target, 1.3° in diameter) and 3 horizontal ellipses (the distractors). “Difficult” distractors subtended 1.6°x1° and “easy” distractors subtended 2.1°x0.5°. On fixed-difficulty search trials, all distractors were either difficult (all-difficult search) or easy (all-easy search). On mixed-difficulty search trials, each display contained one difficult distractor and 2 easy distractors. A black dot (0.1° in diameter) appeared on the left or right side of each shape (0.1° from the outside), with each display containing exactly 2 left-dot and 2 right-dot shapes.

**Figure 1.**
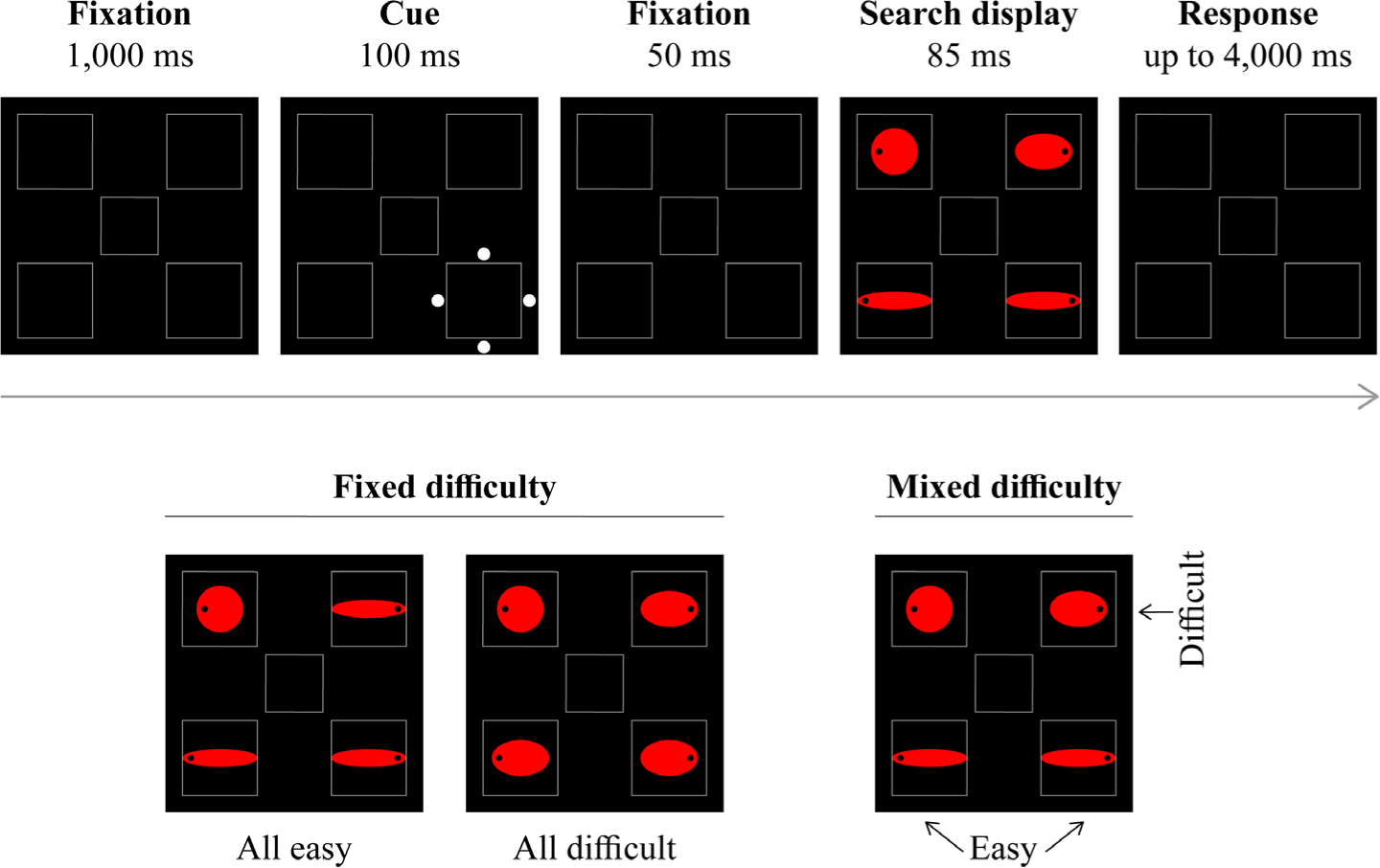
*Upper panel*: Sample sequence of events in Experiment 1. Participants searched for the circle in the target display and reported the side of the black dot inside the target (left or right). This example corresponds to an invalid-cue trial in the mixed-difficulty search condition. The cued distractor is an easy distractor and the response associated with it (right) is incompatible with the response associated with the target (left). *Lower panel*: Sample displays in each search type condition (all-easy, all-difficult, mixed-difficulty). The stimuli are not drawn to scale.

### Procedure

Participants were instructed to search for the circle target and report on which of its sides (left or right) a black dot appeared. They were asked to respond as accurately as possible with no time pressure, and to guess if unsure. They were instructed to press the key Z with their left hands or M with their right hands on the computer keyboard if the dot appeared on the left or on the right, respectively. Each trial began with the fixation display for 1,000ms, followed by the cue display for 100ms and then again by the fixation display for 50ms. Then, the search display appeared for 85ms^4^. A new trial began following the participant’s response or after 4,000ms, whichever came first. Following an incorrect response, participants heard an error beep (225 Hz) for 300ms.

### Design

The experiment consisted of 64 practice trials, followed by 8 blocks of 80 trials each. All-difficult (25%), all-easy-(25%) and mixed-difficulty (50%) search trials were randomly mixed within each block of trials. Conditions of onset-cue location, target location and location of the black dot in each shape (left or right) were equiprobable and randomly mixed within each block of trials. Thus, the cue location was not predictive of the target location.

## Results

The data from three participants were excluded because their mean error rate exceeded the group’s mean by more than two standard deviations (46.6%, 41.9%, and 40.8% vs. M = 21.6%, SD=7.7%). They were replaced by three new participants. Mean accuracy rates are presented in Table 1.

**Table 1.** Mean accuracy rates (in percentage) in Experiment 1 as a function of cue validity (valid vs. invalid) and search type (all-easy, all-difficult and mixed). In the invalid-cue condition, accuracy rates are presented separately for trials in which the cued distractor was compatible vs. incompatible with the target. For the mixed-difficulty condition, either an easy or a difficult distractor could be cued on invalid-cue trials (mixed-easy and mixed-difficult columns, respectively, with valid-cue trials referring to identical trials in the two conditions). The numbers between square brackets represent condition-specific, within-subject 95% confidence intervals (Morey, 2008).

|  | *All-easy* |  | *All-difficult* |  | *Mixed-easy* |  | *Mixed-difficult* |  |
| --- | --- | --- | --- | --- | --- | --- | --- | --- |
| *Valid* | 86.1% | [1.4%] | 75.1% | [1.3%] | 80.7% | [1.0%] | 80.7% | [1.0%] |
| *Invalid* | 86.7% | [1.0%] | 70.2% | [1.2%] | 78.1% | [0.8%] | 74.4% | [1.1%] |
| *Compatible* | 86.4% | [1.1%] | 71.6% | [2.0%] | 74.3% | [1.3%] | 88.6% | [1.3%] |
| *Incompatible* | 86.8% | [1.2%] | 69.9% | [1.4%] | 80.0% | [0.8%] | 67.5% | [1.7%] |

### Effects of search difficulty on valid-cue trials

On valid-cue trials, the effect of search difficulty was significant, F (2, 40) = 18.91, p < .0001, η^2^_p_ = .49, with fewer errors the easier the search (see Table 1). Planned comparisons showed that accuracy was higher on all-easy than on mixed-difficulty trials, F(1, 20) = 9.42, p =.006, η^2^_p_ = .32, and on mixed-difficulty than on all-difficult trials, F(1, 20) = 12.12, p =.002, η^2^_p_ = .38.

### Cue validity effects

Mean cue validity effects on accuracy are presented in Figure 2 (left panel). An analysis of variance (ANOVA) with cue validity (valid vs. invalid) and search difficulty (all-easy, mixed, all-difficult) as factors revealed significant main effects, F (1, 20) = 7.30, p = .01, η^2^_p_ = .27, and F (2, 40) = 53.08, p < 0.0001, η^2^_p_ = 0.73, respectively. The interaction between the two factors was also significant, F (2, 40) = 5.91, p = .006, η^2^_p_ = .23, indicating that the cue validity effect was larger on mixed-difficulty than on all-easy trials, F(1, 20) = 8.78, p = .008, η^2^_p_ = .31, but similar on all-difficult and on mixed-difficulty trials, F < 1. The cue validity effect was significant in the all-difficult and mixed-difficulty conditions, F(1, 20) = 7.78, p = .01, η^2^_p_ = .28 and F(1, 20) = 10.19, p = .005, η^2^_p_ = .34, respectively, and non-significant in the all-easy condition, F<1.

**Figure 2.**
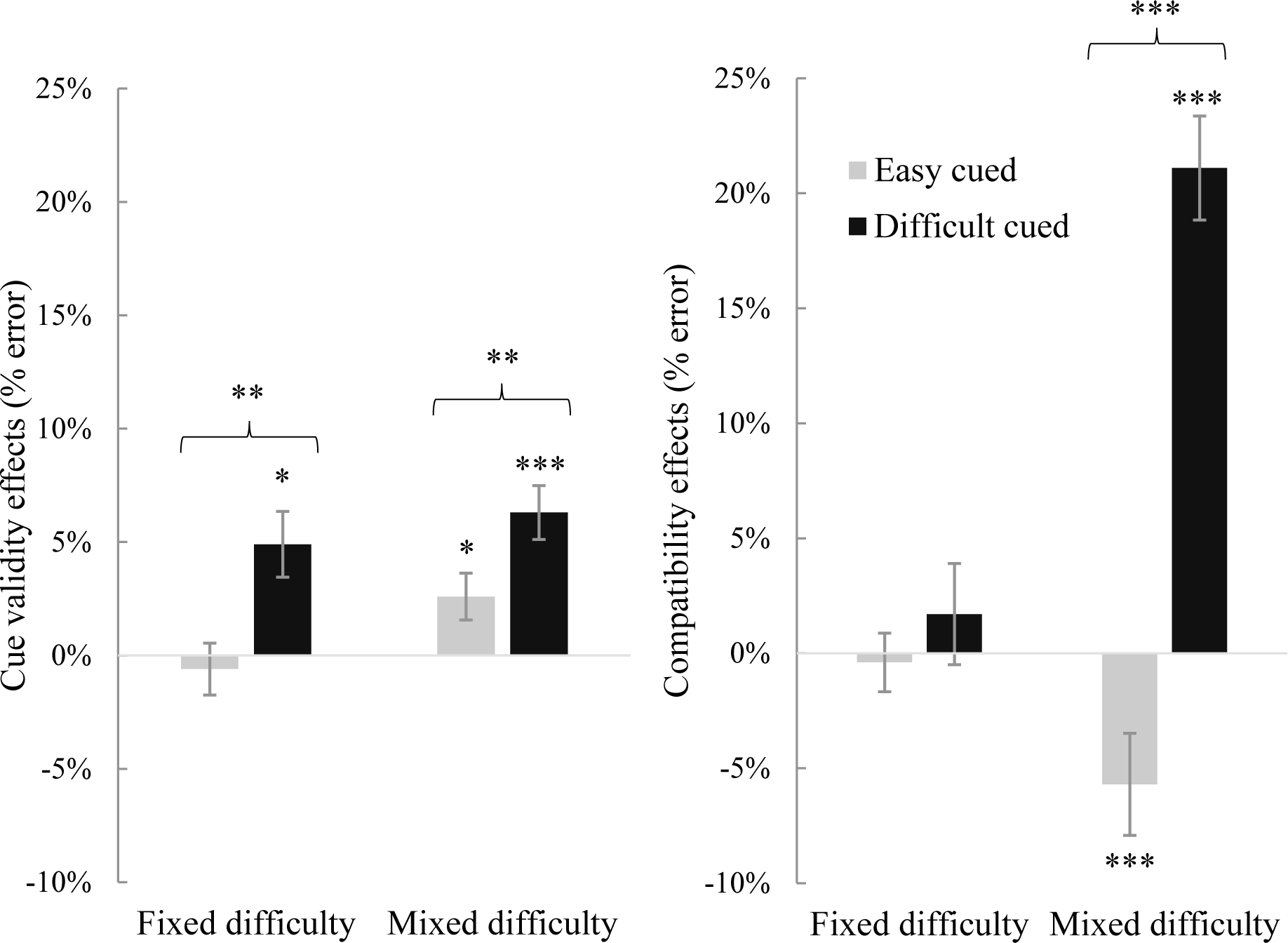
Mean cue effect on target search performance accuracy (in percentage) of cue validity (invalid-cue minus valid-cue, *left panel*) and of cued distractor-target compatibility (incompatible minus compatible, *right panel*) as a function of the search condition: fixed difficulty (all distractors are identical) vs. mixed difficulty (two easy and one difficult distractors appear in the same display) and cued distractor in the invalid-cue condition: easy-cued vs. difficult-cued in Experiment 1. Error bars represent condition-specific, within-subject 95% confidence intervals (Morey, 2008).

Planned comparisons showed that in the mixed-difficulty condition, accuracy was lower when the difficult than when an easy distractor was cued, F(1, 20) = 7.96, p = .01, η^2^_p_ = .29, with poorer performance in both conditions relative to the valid-cue condition, F(1, 20) = 17.84, p=.0004, η^2^_p_ = .47, and F(1, 20) = 4.12, p =.05, η^2^_p_ = .17, respectively.

### Effects of cued distractor compatibility

We conducted a three-way ANOVA on invalid-cue trials with compatibility of the cued distractor (compatible vs. incompatible), cued distractor difficulty (easy vs. difficult) and search type (fixed-vs. mixed) as independent variables (see Figure 2, right panel).

The main effects of compatibility and cued distractor difficulty were significant, F (1, 20) = 17.32, p = .0005, η^2^_p_ = .46 and F (1, 20) = 54.96, p < .0001, η^2^_p_ = .73, respectively, and so were the 2-way interactions between compatibility and cued distractor, F (1, 20) =34.74, p < .0001, η^2^_p_ =.64, between compatibility and search type, F (1, 20) = 12.96, p =.002, η^2^_p_= .39 and between cued distractor difficulty and search type, F(1, 20) = 79.1, p <.0001, η^2^ = .80. These were modulated by a significant three-way interaction, F (1, 20) = 58.66, p<.0001, η^2^_p_ = .75.

In order to clarify this interaction, we conducted separate analyses for each search type condition. When distractor difficulty was fixed, there was no compatibility effect, irrespective of whether the distractors were easy or difficult, both Fs < 1. When distractor difficulty was mixed, the compatibility effect was larger when the cued distractor was difficult than when it was easy, F (1, 20) = 63.41, p < .0001, η^2^_p_ = .76. Planned comparisons revealed that when the difficult distractor was cued, accuracy was higher when this distractor was compatible than when it was incompatible, F(1, 20) = 74.17, p < .0001, η^2^_p_ = 0.79, and the compatibility effect was significant but in the opposite direction when an easy distractor was cued, F(1, 20) = 14.18, p = .0001, η^2^_p_ = .41.

### Effects of difficult-distractor compatibility

Mean accuracy rates are presented in Table 2. An ANOVA with compatibility between the difficult distractor and the target (compatible vs. incompatible) and cued object (target, easy distractor, difficult distractor) as factors revealed significant main effects, F(1, 20) =85.76, p < .0001, η^*2*^_*p*_ = .81 and F(2, 40) = 9.54, p = .0004, η^*2*^_*p*_ = .32, respectively. The interaction between the two factors was also significant, F(2, 40) = 4.69, p = .01, η^*2*^_*p*_ = .19, indicating that the effect of the compatibility between the difficult distractor and the target was larger when the difficult distractor itself was cued than when the easy distractor was cued, F(1, 20) = 5.20, p = .03, η^*2*^_*p*_ = .21 and was similar irrespective of whether the target or an easy distractor was cued, F < 1. Crucially, paired comparisons showed that the effect of the compatibility between the difficult distractor and the target was significant in all three conditions, all ps <.0001, all η^*2*^_*p*_ > .64.

**Table 2.** Mean accuracy rates (in percentage) in the mixed-difficulty condition of Experiment 1, as a function of cued object (target, easy distractor, difficult distractor) and compatibility between the difficult distractor and the target (compatible vs. incompatible). The numbers between square brackets represent condition-specific, within-subject 95% confidence intervals (Morey, 2008).

|  | *Target cued* |  | *Easy distractor cued* |  | *Difficult distractor cued* |  |
| --- | --- | --- | --- | --- | --- | --- |
| *Compatible* | 90.7% | [1.7%] | 87.8% | [1.3%] | 88.6% | [1.3%] |
| *Incompatible* | 75.9% | [1.5%] | 73.5% | [1.1%] | 67.5% | [1.7%] |

## Discussion

Overall, the data closely conformed to the predictions of PAF and provided a clear response to the main question addressed by Experiment 1. Although search displays were presented too briefly for participants to engage in verification processes after locating the target, accuracy when the target was cued depended on how similar the distractors were to the target. While this finding can be readily explained by PAF, it is incompatible with the standard interpretation of the spatial cueing effect (e.g., Folk et al., 1992) and with the dwelling hypothesis (Gaspelin et al., 2016). These both predict that performance on valid-cue trials should be similar irrespective of search difficulty because the spatial cueing effect reflects the fact that attention resides at the cued location.

One may argue that the critical distractor-similarity effect on valid-cue trials might reflect the performance of a few participants who were able to locate the target and engage in verification processes, within 85ms. If so, the participants who were most accurate on the search task should be the ones who had time to verify that the cued object was indeed the target. However, this alternative account is unlikely because the finding was replicated when only the 50% of participants who were least accurate on valid-cue trials (M = 74%) were considered, F(2, 20) = 10.66, p = .0007, η^*2*^_*p*_= .52.

We replicated the other main findings reported by Lamy et al. (2018), which are equally compatible with the dwelling hypothesis and with PAF. The cue validity effect increased as overall target-distractor similarity increased. In the mixed-difficulty condition, accuracy was higher when an easy than when a difficult distractor was cued, and the compatibility between the difficult distractor and the target strongly affected performance, both when this distractor was cued and when the easy distractor was cued.

However, we also found that the compatibility between the difficult distractor and the target affected performance even when the target was cued. This result is not predicted by the dwelling hypothesis because the difficult distractor should only rarely become the focus of attention when the target is cued. By contrast, it is easily accommodated by PAF: when competition is high, an uncued difficult distractor may be more activated than the cued target on some trials (at least when the cue does not match the search template). On such trials, attention is shifted to the difficult distractor, hence the compatibility effects.

There were three exceptions. First, the cue validity effect was not significant when all the distractors were easy. It is possible that with a brief relative to a long exposure of the search display, participants maintain a stronger attentional set. In that case, the target should win the competition equally fast, irrespective of whether or not it is cued, hence the null effect. In any event, since the qualitative pattern was replicated – namely, the cue validity effect was smallest in that condition - this finding does not challenge any of our conclusions.

Second, in the mixed-difficulty condition, when the easy distractor was cued, the compatibility effect was negative: accuracy was higher when the easy (cued) distractor was incompatible with the target than when it was compatible. This finding is most likely to result from an idiosyncratic feature of the present design. There were two objects with a left-side dot and two with a right-side dot. Thus, when the cued easy distractor was compatible with the target, the difficult distractor was necessarily incompatible with it. Since we also found that the compatibility of the difficult distractor with the target strongly modulated performance even when this distractor was not cued, the negative compatibility effect is likely to index processing of the difficult (uncued) distractor rather than processing of the easy cued distractor (see Lamy, Darnell, Levi & Bublil, 2019 for a similar finding).

Finally, no compatibility effect was observed in the all-difficult condition. It is difficult for the dwelling hypothesis to accommodate this finding: unlike when viewing time is unlimited, there should not be enough time to disengage from the cued difficult distractor’s location when search display exposure is limited to 85ms. According to PAF, however, the large cue-validity effect in the all-difficult condition does not necessarily indicate that attention was directed to the cued location: instead, it indicates that the competition in favor of the target is resolved faster when the target is cued than when it is not, but when the competition is strong and the cue weight relatively small, as was the case here in the all-difficult condition, attention may often accrue to an uncued difficult distractor, and the compatibility effect from the cued distractor should therefore be diluted.

### Experiment 2

The results of Experiment 1 refute the argument according to which RTs become slower as search difficulty increases on valid-cue trials (Gaspelin et al., 2016; Lamy et al., 2018) because participants engage in verification processes that take more time, the more similar the distractors are to the target. However, these results do not address the second alternative account raised in the introduction, namely, that the search-difficulty gradient on valid-cue trials might emanate from trials in which the cue did not capture attention.

In Experiment 2, we tested this possibility by conducting new analyses on the data from Lamy et al.’s (2018) first experiment. This experiment was identical to Experiment 1 except that the search display was presented until response and responses were speeded. According to the standard interpretation of cue validity effects, on valid-cued trials in which the cue does capture attention, attention is already focused at the target location when the search display appears. It follows that these trials should be the fastest trials and crucially, they should be equally fast in the three conditions of search difficulty.

We tested this prediction by plotting the distribution of the trials in which the target appeared at the same location as the cue. To do that, we used a vincentization procedure (Ratcliff, 1979): quantiles of RT distributions were computed for each participant, each summarizing 10% of the cumulative RT distribution, and were then averaged to produce the group distribution (Rouder & Speckman, 2004). This nonparametric procedure was applied on valid-cue trials separately for the all-easy, mixed-difficulty and all-difficult conditions. As is clear from Figure 3, the effect of search difficulty was present across the RT distribution. In particular, it was already large and significant for the 10% fastest trials, F (2, 46) = 47.59, p < .0001, η^*2*^ _*p*_ = .67. Thus, for the partial attentional capture account to explain these data, one would have to assume that the cue captured attention on much less than 10% of the trials. Such a conjecture would considerably weaken the relevance of cue validity effects as indicators of involuntary capture of attention. Finally, note that the finding that the critical effect was present on the fastest trials also invalidates the claim that it reflects verification processes: the fastest trials should be those in which participants did not engage in such verification.

**Figure 3.**
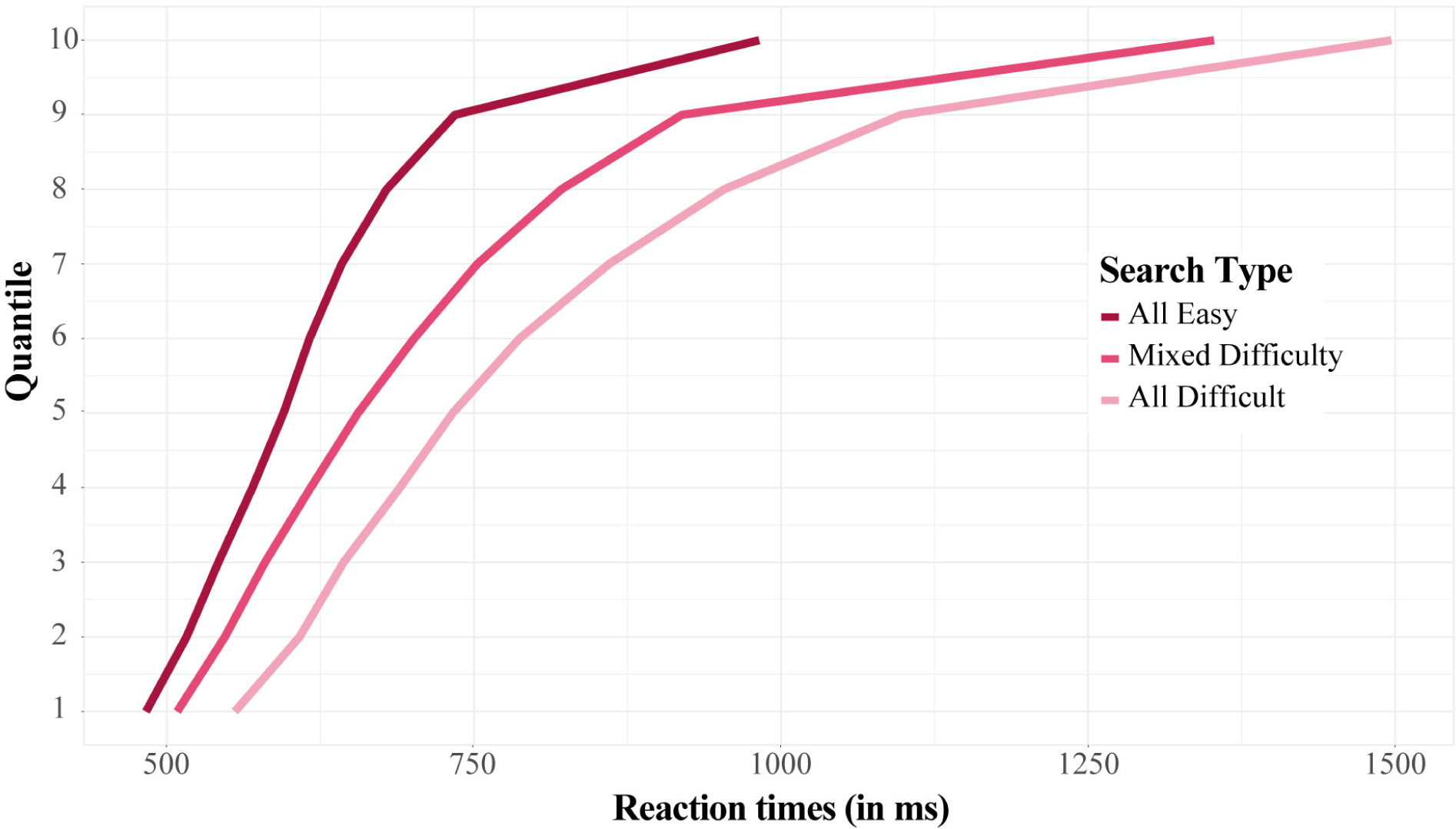
Vincentized reaction time distribution (quantile means) in all-easy, mixed difficulty and all-difficult search conditions, on valid-cue trials in which the target appeared at the cued location in experiment 2.

Taken together with the findings from Experiment 1, the outcomes of the new analyses presented here strongly support PAF’s interpretation. Attention is not shifted to the cued location immediately following its onset. Instead, attention is allocated in the search display to the location that has accumulated the highest priority, and such allocation occurs faster the more discriminable the distractors are from the target. Therefore, search difficulty affects performance even when the target is validly cued.

### Experiment 3

So far, we have suggested that while cue validity effects are not necessarily diagnostic of attentional allocation, effects of distractor-target compatibility indicate that attention was shifted to the location of the distractor (provided that discriminating the response feature requires focused attention). Yet, compatibility effects only provide a measure of whether attention was allocated to a specific location and gives no indication as to how much priority accrued to the other locations. In Experiment 3, we sought a measure of the distribution of spatial attention across the search display.

Previous studies have used the dot-probe paradigm in order to obtain a snapshot of the distribution of attention during visual search (e.g., Kim & Cave 1995; Lamy, Tsal & Egeth 2003; Watson & Humphreys 2000). In a typical dot-probe study, the search display is immediately followed, on a portion of the trials, by a probe that is equally likely to appear at each of the previously filled locations. Participants are required to make a non-speeded response to the target on each trial, but to first press a key as fast as possible whenever they detect a probe. The crux of this paradigm is that, because participants do not know on any given trial whether they will have to respond to the target (probe-absent trials) or to the probe (probe-present trials), they search for the target on both types of trials. Thus, how fast and accurate participants are to respond to the probe is thought to index how much attention was allocated to the object that occupied the probe location while they searched the display for the target.

A prominent advantage of the dot-probe method is that it allows one to assess the relative amount of attention accruing not only to the target but also to non-target locations. This feature is particularly useful in the present context because it should help us uncover what modulates attentional activation at each location: (a) whether or not it was cued, (b) how similar the object at that location is to the target and (c) how strong the competition is.

In Experiment 3, we thus combined the spatial cueing and dot-probe paradigms in order to further test the priority accumulation framework against the dwelling hypothesis. It was similar to Experiment 1, with two notable changes. A probe immediately followed the offset of the search display on half of the trials and participants were required to respond as fast and accurately as possible to its presence before making their non-speeded response to the target. In addition, only the mixed difficulty condition was included.

According to the dwelling hypothesis, after the cue appears, attention is allocated to the cue location, remains at that location until the search display appears and is then disengaged from that location more or less slowly, depending on how similar the distractor appearing at the cued location is to the target. Thus, performance at detecting a probe flashed very shortly after the search display should be invariably best when the probe appears at the cued location, irrespective of what object is cued.

According to PAF, attentional priority at a given location is determined by the characteristics of the events that occur at that location across time (i.e., events such as the cue and the search display objects), and in particular, by how salient and how similar they are to the target. Thus, what object is cued also determines how strong the competition is and how long it should take to resolve: for instance, the target should suffer from more competition when the difficult distractor is cued than when an easy distractor is cued.

We conducted four analyses in order to test these predictions against each other. First, we analyzed cue-validity and compatibility effects on target discrimination performance when the probe was absent. We expected to replicate all the results observed in the mixed-difficulty condition of Experiment 1. This replication was crucial to ensure that we could use probe-detection performance on probe-present trials as a proxy of the distribution of attention during search in the mixed-difficulty condition.

Then, we examined probe-detection performance when the probe was present. We asked whether when an easy distractor is cued, probe detection is better at the location of that distractor, as predicted by the dwelling hypothesis, or, at the location of the target, as predicted by PAF. The latter result would provide important converging evidence for our interpretation of a finding central to PAF: the spatial cueing effect observed in the absence of a compatibility effect in the mixed-difficulty condition (Exp.1, see also Lamy et al., 2018). We suggested that in that condition, attention was not shifted to the cue, because the easy distractor was very dissimilar to the target and the extra activation provided by the cue was not enough to allow its location to win the competition: the cue validity effect only reflected the fact that the competition was resolved slower when the easy distractor was cued than when the target was cued.

Next, we tested PAF’s prediction that probe detection performance should be improved both (a) when the probe appears at the location of the cue and (b) the more similar to the target the probed object is.

Finally, we tested PAF’s claim that what object is cued determines how strong the priority advantage of the winning object is. Specifically, PAF predicts that the extent to which the identity of the probed object (target, easy distractor or difficult distractor) modulates probe-detection performance should be maximal when the target is cued (with fast resolution of the competition), intermediate when an easy distractor is cued and minimal when the difficult distractor is cued (with slow and inconsistent resolution of the competition).

## Methods

### Participants

Twenty-one undergraduate students (11 female, age M = 25.73, SD =3.76) participated for course credit. Participants reported normal or corrected-to-normal visual acuity.

### Apparatus, stimuli, procedure and design

A sample sequence of events is presented in Figure 4. The apparatus, stimuli, procedure and design were similar to those of Experiment 1, except for the following changes. Only the mixed-difficulty condition was included. All shapes in the search display were ellipses, subtending 1.4°×1.21° for the target, 1.6°×1.06° for the high-similarity (difficult) distractor and 2°×0.84° for the two low-similarity (easy) distractors. The target was defined as the object that most resembled a circle^5^. The probe was a small blue X (RGB=0, 0, 255) subtending 0.3° inside.

**Figure 4.**
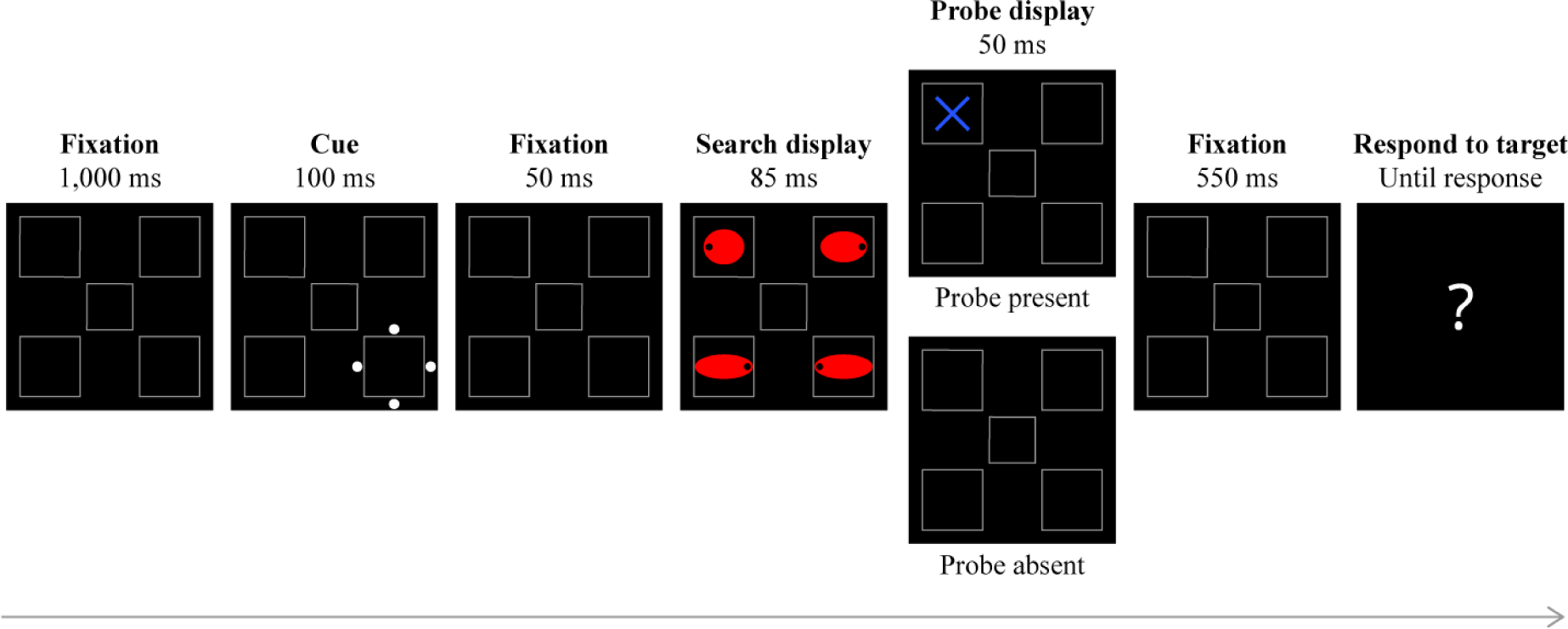
Sample sequence of events in Experiment 2. It was similar to Experiment 1 except that a probe-detection task was added. Prior to responding to the target, participants had to press a key within 600ms when the probe was present (probe-present trials, 50% of all trials) and to refrain from pressing this key when the probe was absent (probe-absent trials, 50% of all trials). The stimuli are not drawn to scale.

On 50% of the trials (probe-present trials), a probe appeared for 50ms at the center of one of the four possible stimulus locations, immediately following the search display offset. Then, the fixation display appeared for 550-ms, followed by a question mark. On the remaining 50% of the trials (probe-absent trials), the fixation display replaced the probe display and thus appeared for 600ms. Participants were instructed to report the presence of the probe, as quickly as possible within the 600-ms response window, by pressing “0” on the numerical keypad using their right-hand index fingers, and to respond to the target after the question mark appeared (by pressing “Z” for left, and “X” for right, using their left hands). The error beep was sounded if a participant’s response was wrong for either the probe or the search task.

The experiment started with two 15-trial practice blocks. In the first practice block, participants were required to respond only to the target. The second practice block was similar to the experimental blocks: the probe appeared on 50% of the trials and participants responded first to the probe and then to the target. There were nine experimental blocks of 96 trials each. On probe-present trials, the probe was equally likely to appear at each of the four possible locations. All conditions were randomly mixed within each block of trials.

## Results

The data from five participants were excluded because their performance deviated from the group’s mean by more than 2 standard deviations: % of misses on the probe task (one participant, 79% vs. M = 14.6%, SD =9.1%), % of false alarms on the probe task (three participants, 40.6%, 44.5% and 35.7% vs. M = 7.8%, SD = 6.3%) and % of errors on the search task on probe-absent trials (one participant, 48.2% vs. M = 25.8%, SD = 8.9%). They were replaced by five new participants. No participant met the exclusion criterion on % errors on the search task when the probe was present (M=33.4%, SD=10.1%).

### Search task performance

Performance on the search task was significantly impaired when participants also had to respond to the probe (probe-present trials), F (1, 20) = 34.89, p < .0001. To verify that we replicated the findings observed in the mixed-difficulty condition of Experiment 1, we first analyzed performance on the search task when the probe was absent and participants correctly refrained from hitting the zero number-key (92.2% of the trials). Mean spatial cueing effects and mean compatibility effects are presented in Figure 5. Mean accuracy rates are presented in Table 3.

**Table 3.** Mean accuracy rates (in percentage) on the target search task in Experiment 3 as a function of cue validity (valid vs. invalid). In the invalid-cue condition, the accuracy rates are presented separately as a function of whether the cued distractor was easy or difficult and of whether it was compatible or incompatible with the target. The numbers between square brackets represent condition-specific, within-subject 95% confidence intervals (Morey, 2008).

|  |  |  |  |  |
| --- | --- | --- | --- | --- |
| *Valid (Target cued)* | 82.1% | [1.8%] |  |  |
|  | *Easy cued* |  | *Difficult cued* |  |
| *Invalid* | 73.8% | [1.0%] | 69.3% | [1.7%] |
| *Compatible* | 68.2% | [1.1%] | 90.7% | [1.9%] |
| *Incompatible* | 76.3% | [0.8%] | 59.7% | [2.2%] |

**Figure 5.**
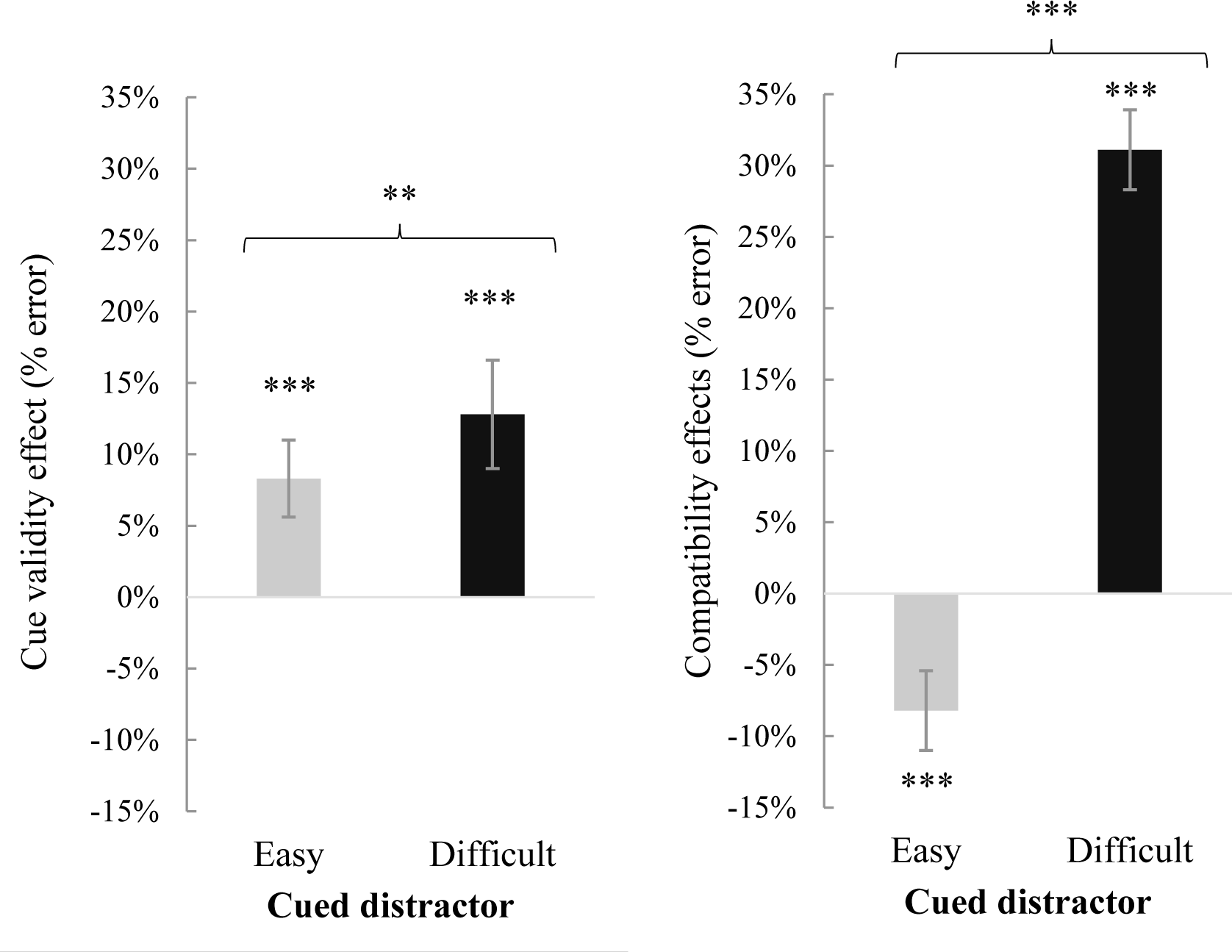
Mean effect on target search performance accuracy (in percentage) of cue validity (invalid-cue minus valid-cue, *left panel*) and of cued distractor-target compatibility (incompatible minus compatible, *right panel*) as a function of cued distractor for the invalid-cue condition (easy vs. difficult) in Experiment 3. Error bars represent condition-specific, within-subject 95% confidence intervals (Morey, 2008).

### Cue validity effects

Planned comparisons revealed significant cue validity effects, with higher accuracy on valid-relative to invalid-cue trials, both when the cued distractor was easy, F(1, 20) = 19.03, p = .0003, η^*2*^ _*p*_ = .49, and when it was difficult, F(1, 20) = 21.83, p = .0001, η^*2*^_*p*_ = .52. Accuracy was higher when the easy than when the difficult distractor was cued, F (1, 20) = 7.33, p = .01, η^*2*^_*p*_ = .27.

### Effects of cued distractor compatibility

An ANOVA on invalidly cued trials, with cued object (easy vs. difficult) and response compatibility (compatible vs. incompatible) as factors revealed significant main effects of cued object, F (1, 20) = 5.67, p = .03, η^*2*^_*p*_ = .22, and compatibility, F (1, 20) = 35.98, p < .0001, η^*2*^_*p*_ = .64, as well as a significant interaction between the two factors, F (1, 20) = 97.2, p < .0001, η^*2*^_*p*_ = .83. Paired comparisons revealed that compatible trials were more accurate than incompatible trials when the cued distractor was difficult, F (1, 20) = 72.4, p = .0001, η^*2*^_*p*_ = .78, and that the effect was in the opposite direction when the cued distractor was easy, F (1, 20) = 35.8, p = .0001, η^*2*^_*p*_ = .64.

### Effects of difficult-distractor compatibility

Next, we examined the effect of the compatibility between the difficult distractor and the target (compatible vs. incompatible) as a function of which object was cued (target, difficult distractor, easy distractor). Mean accuracy rates are presented in Table 4. Both main effects were significant, F (1, 20) = 121.3, p < .0001, η^*2*^_*p*_ = .85 and F (2, 40) = 18.29, p < .0001, η^*2*^_*p*_ = .48, respectively. The significant interaction between the two factors, F (2, 40) = 9.26, p = .0003, η^*2*^ = .33, indicated that the compatibility effect was larger when the difficult distractor was cued than when either the target or the easy distractor was cued, F (1, 20) = 11.84, p = .003, η^*2*^_*p*_ = .37 and F(1, 20) = 11.52 p = .003, η^*2*^_*p*_ = .37, respectively. Crucially, paired comparisons showed that the effect of the compatibility between the difficult distractor and the target was significant in all three conditions, all ps <.0001, all η^*2*^_*p*_ > .59.

**Table 4.** Mean accuracy rates (in percentage) on the target search task in the mixed-difficulty condition of Experiment 3, as a function of cued object (target, easy distractor, difficult distractor) and compatibility between the difficult distractor and the target (compatible vs. incompatible). The numbers between square brackets represent condition-specific, within-subject 95% confidence intervals (Morey, 2008).

|  | *Target cued* |  | *Easy cued* |  | *Difficult cued* |  |
| --- | --- | --- | --- | --- | --- | --- |
| *Compatible* | 91.8 % | [1.2%] | 86.3% | [1.6%] | 90.7% | [1.7%] |
| *Incompatible* | 77.0% | [2.5%] | 67.3% | [1.3%] | 59.7% | [2.7%] |

### Probe-task performance

Probe-absent trials were excluded from all the following analyses. Preliminary analyses on probe-present trials indicated that there was no speed-accuracy trade-off, with a strong negative correlation between probe RTs and accuracy. We could thus integrate the two measures into a single measure, the Inverse Efficiency Score (IES; Townsend & Ashby, 1978), which is obtained by dividing the mean RT by the mean accuracy rate for each participant for each condition. It has been suggested that this measure can be useful to detect small effects, provided that the speed and accuracy data are also inspected (e.g., Vandierendonck, 2017). As is clear from a comparison of the probe-detection mean RTs, accuracy and IES scores presented in Table 5, all the relevant effects (i.e., the ranking of the scores within each row of Table 5) followed the same pattern on RTs and accuracy.

**Table 5.** Mean reaction times (in milliseconds), accuracy rates (in percentage) and Inverse Efficiency Scores (IES, in milliseconds) on the probe detection task in Experiment 3, as a function of cued object (target, easy distractor or difficult distractor) and probed object (target, easy distractor or difficult distractor). The numbers between square brackets represent condition-specific, within-subject 95% confidence intervals (Morey, 2008). Note that the mean IES scores are different from the ratio of the mean RT by the mean accuracy score because this ratio was calculated separately for each subject and then averaged.

|  | *Probe on target* |  | *Probe on easy* |  | *Probe on difficult* |  |
| --- | --- | --- | --- | --- | --- | --- |
| RTs |  |  |  |  |  |  |
| *Target cued* | 418 | [3] | 427 | [3] | 429 | [5] |
| *Easy cued* | 415 | [2] | 420 | [3] | 421 | [4] |
| *Difficult cued* | 425 | [3] | 426 | [3] | 417 | [3] |
| Accuracy |  |  |  |  |  |  |
| *Target cued* | 89.6% | [1.4%] | 84.1% | [1.3%] | 86.7% | [1.8%] |
| *Easy cued* | 86.8% | [0.8%] | 85.7% | [1.0%] | 84.5% | [1.0%] |
| *Difficult cued* | 85.8% | [1.8%] | 84.2% | [1.3%] | 86.0% | [1.1%] |
| IES |  |  |  |  |  |  |
| *Target cued* | 473 | [9] | 517 | [8] | 501 | [13] |
| *Easy cued* | 484 | [5] | 500 | [8] | 508 | [8] |
| *Difficult cued* | 511 | [15] | 519 | [10] | 495 | [8] |

A planned comparison indicated that when the easy distractor was cued, probe detection was better when the target’s location than when the easy (cued) distractor’s location was probed, F(1, 20) = 4.70, p = .04, η^2^_p_= .19. Thus, the critical prediction of PAF was confirmed.

Next, we assessed how for each location, the occurrence of the cue and the similarity to the target of the object at that location, affected the amount of attention at that location in the search display. To do that, we conducted an ANOVA with cue-probe location (same vs. different) and probed object (target, easy distractor or difficult distractor) as within-subject factors.

The main effect of cue-probe location was significant, F (1, 20) = 6.27, p = .02, η^*2*^_*p*_ = .24, with better performance when the probe appeared at the cued than at an uncued location. The main effect of probed object was also significant, F(2, 40) = 7.60, p = .002, η^*2*^_*p*_ = .28, indicating that performance was better when the probe appeared at the location of the target than at the difficult distractor’s location, F(1, 20) = 9.25, p = .006, η^*2*^_*p*_ = .32, and was similar when the difficult vs. an easy distractor’s location was probed, F(1, 20) = 1.11, p = .3, η^*2*^_*p*_ = .01. There was no significant interaction, F < 1. These results confirm that probe detection performance was improved both when the probe appeared at the location of the cue and the more similar the probed object was to the target - although the numerical difference between difficult- and easy-distractor probed trials (500 vs. 508ms, respectively) did not reach significance.

Finally, we measured the impact of the competition on the attentional benefit accruing to the target’s location relative to other locations. To do that, we conducted an ANOVA with cued object (target, easy distractor or difficult distractor) and probed object (target, easy distractor or difficult distractor) excluding trials in which an easy distractor was probed and the other easy distractor was cued. This exclusion was necessary in order to perform a balanced comparison between the target and difficult distractor on the one hand, of which there was only one, and the easy distractor on the other hand, of which there were two.

The main effect of probed object was significant, F(2, 40) = 4.55, p = .02, η^*2*^_*p*_ = .19, indicating that probe detection was better when the target than when either the difficult or an easy distractor was probed, F(1, 20) = 3.43, p = .04, η^*2*^_*p*_ = .15 and F(1, 20) = 9.05, p = .0006, η^*2*^_*p*_ = .31, respectively, with no difference between the latter two conditions, F < 1. The interaction between cued object and probed object was marginally significant, F (4, 80) = 2.46, p = 0.05, η2p = .11. Planned comparisons revealed that the effect of probed object was highly significant when the target was cued, F(2, 40) = 5.69, p = 0.007, η^*2*^_*p*_ = .22, marginally significant when an easy distractor was cued, F(2, 40) = 3.14, p = 0.05, η^*2*^_*p*_ = .14, and non-significant when the difficult distractor was cued, F(2, 40) = 1.27, p = 0.29, η^*2*^_*p*_ = .06. These results confirm that the easier it was to resolve the competition (which was determined here by what object is cued), the larger the benefit when the probe appeared at the target’s location relative to the difficult or easy distractors’ locations.

**Figure 6.**
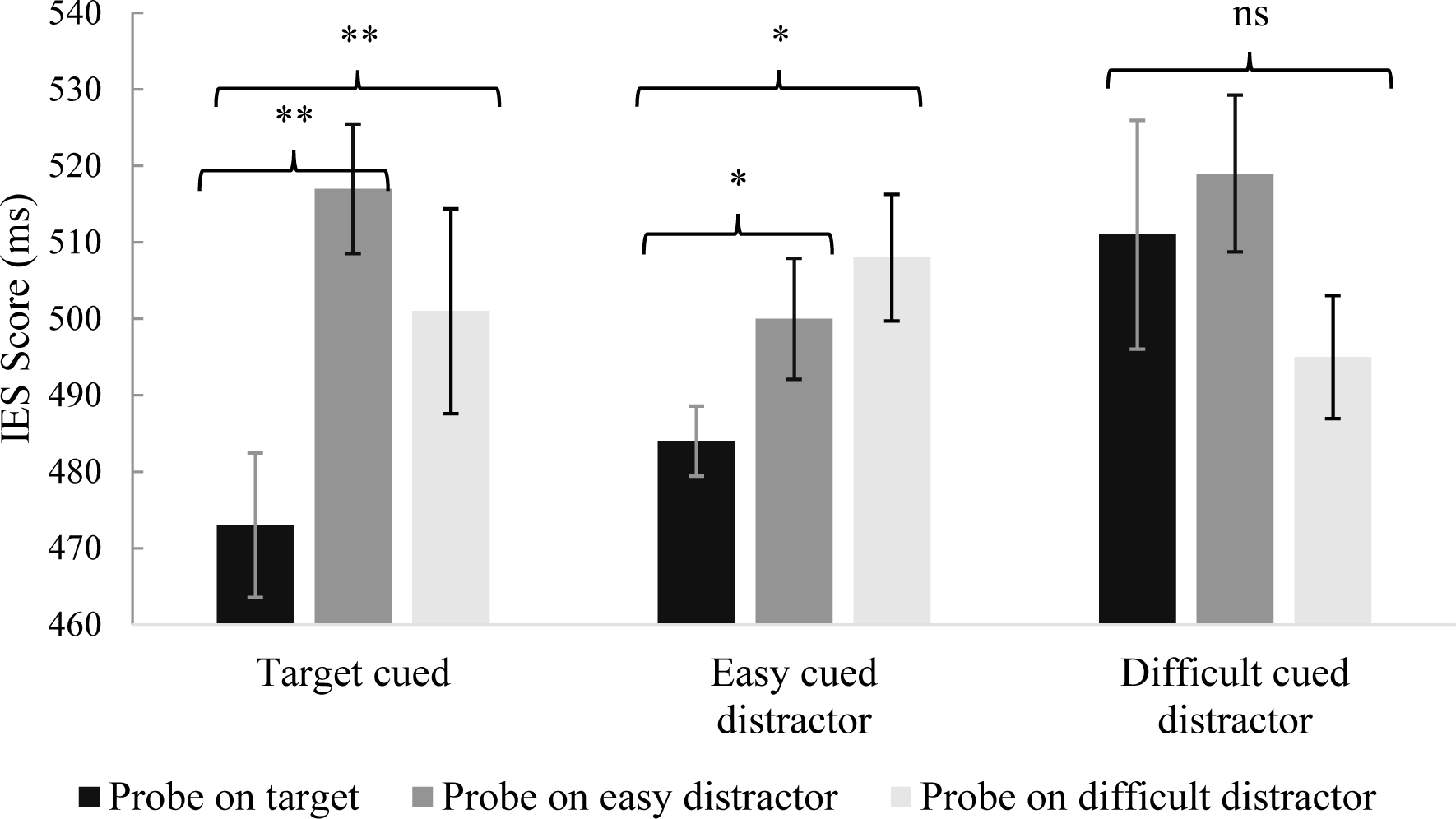
Mean Inverse Efficiency Scores (IES, in milliseconds) in Experiment 3, as a function of cued object (target, easy distractor or difficult distractor) and probed object (target, easy distractor or difficult distractor). Error bars represent condition-specific, within-subject 95% confidence intervals (Morey, 2008).

## Discussion

On the search task, the results of Experiment 3 replicated all the findings observed in the mixed-difficulty condition of Experiment 1: on probe-absent trials, search was most accurate when the target was cued and, crucially, search was also more accurate when the easy distractor was cued than when the difficult distractor was cued. In addition, the compatibility of the response feature associated with the difficult distractor with the response feature associated with the target affected performance, wherever the cue appeared, but more strongly so when the cue appeared at the location of the difficult distractor. Finally, when an easy distractor was cued, we again observed a negative effect of the compatibility between this distractor and the target.

The findings for the probe detection task further supported PAF’s predictions. First, when an easy distractor was cued, performance was higher when the probe appeared at the location of the target than when it appeared at the cued location. This finding is incompatible with the notion that the spatial cueing effect indicates that attention is shifted to the cue location (Folk et al., 1992; Gaspelin et al., 2016; Posner et al., 1980; Theeuwes, 2010). By contrast, it is in line with PAF’s prediction that when an easy distractor is cued (with a “weak” cue), attention is typically not shifted to its location: the target wins the competition and the spatial cueing effect on the search task only indicates that the resolution of the competition is easier when the target is cued than when an easy distractor is cued. Alternative accounts are presented in the General Discussion. Second, probe detection performance was modulated both by whether the probe appeared at a cued or uncued location and by the similarity of the probe object to the target. Finally, the priority advantage of the target relative to the difficult and easy distractors was modulated by what object was cued (target, difficult distractor or easy distractor) indicating that this relative priority advantage determined how easily the competition was resolved.

## General Discussion

Spatial cueing effects have shaped attention theories, based on the premise that they provide a reliable indicator of attentional allocation. Here, we challenge the canonical interpretation, according to which finding spatial cueing effects necessarily entails that attention was shifted to the cued location: we provide evidence showing that spatial cueing effects can be observed even when attention was never directed to the cue. We suggest a new model of attentional allocation, the Priority Accumulation Framework (PAF, see also Gabbay et al., 2019; Lamy et al., 2018) that accounts for many inconsistent findings in the attentional capture literature.

The main idea underlying PAF is that attentional priority weights accumulate at each location of the visual field across time, as a function of the events that occur at each location. This accumulation process terminates when the search context triggers selection, and attention is shifted to the highest-priority location. Crucially, the time it takes for the winner to prevail is shorter the larger the winner’s leading edge. Thus, according to PAF, where a cue appears can bias competition in favor of the cued location (and yield a spatial cueing effect), but does not always determine which location is the winner (i.e., the locus of the first attentional shift).

The Priority Accumulation Framework further posits that the saliency of the events occurring at a given location and the extent to which these events match the observer’s goals (as well as noise) determine how much attentional priority accrues to this location. Thus, a cue’s relative contribution to a location’s attentional priority weight is modest when this cue does not share the target’s defining property and when distractors are very different from the target (which explains findings supporting the contingent-capture account, e.g., Folk et al., 1992).

### Summary of the study’s novel findings

According to the standard interpretation of spatial cueing effects, faster RTs on valid-relative to invalid-cue trials indicate that attention is still focused at the cued location when the search display appears. The target is therefore responded to quickly because there is no need to search any further. According to this interpretation, performance on valid-cue trials should be independent of the similarity between the target and the distractors that surround it in the search array. The Priority Accumulation Framework makes the opposite prediction: RTs on valid-cue trials should be faster when the discrimination between the target and distractors is easy than when it is difficult, because the competition should be resolved faster in the former case. Previous findings (Lamy et al., 2018, see also Gaspelin et al., 2016 and footnote 1) supported this prediction. However, they were open to alternative interpretations: the observed search difficulty gradient might (1) reflect post-attentional verification processes, or (2) result from trials in which of the cue failed to capture attention. In Experiments 1 and 2, we refuted these possibilities: we replicated the critical finding with short search display exposure times that prevented verification processes (Experiment 1) and showed that the search difficulty gradient was strong even for the fastest trials, which should be those in which the cue captured attention.

In Experiment 3, we tested novel predictions of PAF by combining the spatial cueing paradigm with a dot-probe paradigm that provided a proxy of the distribution of attention across the search display. In particular, we predicted that when the cue appears at the location of an easy distractor, probe detection performance should be better at the target’s than at the cue’s location (based on our claim that attention is not shifted to the cued location in that condition). In addition, we predicted that probe-detection performance should be best when the probe appears at the target location, but that this advantage should depend on how fast the resolution of the competition is (maximal when the target is cued, intermediate when an easy distractor is cued, and minimal or null when the difficult distractor is cued). By contrast, according to the standard rationale underlying spatial cueing experiments, if a cue yields spatial cueing effects, a probe displayed shortly after the search display should be responded to fastest when it appears at the cued location, irrespective of what object appears at that location. The results confirmed all PAF’s predictions.

### Alternative accounts for the findings

Possible alternative accounts for the present findings hinge on the claim that using an 85-ms search display duration (in Experiments 1 and 3) may not have achieved the intended goals.

In Experiment 1, our main objective was to prevent participants from engaging in verification processes after locating the target by shortening display exposure: as such verification processes should be lengthier when the distractors are similar to the target than when they are dissimilar, they would explain why accuracy on valid-cue trials was poorer the lower target-distractor discriminability was.

It is highly unlikely that the 85-ms display duration used here was long enough to allow such verification processes. When accuracy is far below ceiling, participants have no incentive to shift attention from the target to verify that it is indeed the target: in order to maximize their performance, they are more likely to use the search display time to process the target. In Experiment 1, overall accuracy on valid-cue trials was clearly below ceiling (M=81%) and the critical finding was replicated even when only the data from the participants who were least accurate on valid-cue trials were analyzed (M=74%).

In Experiment 3, our selection of the search display exposure duration and search-probe SOA was guided by two objectives: (1) to maintain the overall accuracy rate on the search task above chance but below ceiling (which was achieved with an 85-ms search display exposure in Experiment 1) and (2) to take a snapshot of the distribution of attention shortly after the search display was presented in order to catch the first shift of attention (which is why we had the probe appear immediately after the search display offset).

One may argue that during the 85-ms search display presentation, participants had time to shift their attention away from the cued location when the easy distractor was cued. According to this interpretation, (1) dwelling was too brief to allow processing of the cued easy distractor’s response feature, hence the absence of a compatibility effect, and (2) when the probe was flashed, attention was most often already focused on the target, hence the better probe detection performance at the target than at the cued location. By contrast, (3) when the difficult distractor was cued, disengagement took longer and attention was redirected to the target only on a portion of the trials, hence the significant compatibility effect and the null effect of the probed object on probe detection.

Two findings argue against this account. First, if attention was shifted to the target location within 85ms when an easy distractor was cued, performance on search trials should be similar when the target was cued and when an easy distractor was cued – yet, this was clearly not the case. Second, it is not clear why compatibility effects from the difficult distractor should be observed on the search task when the target was cued. According to the dwelling hypothesis, attention should accrue to the target location on most trials when this location is cued and no compatibility effects are therefore expected. According to PAF, the difficult distractor may win the competition on a portion of the (valid-cue) trials, hence the observed compatibility effect. In addition, this compatibility effect should be smaller the smaller the probability that the difficult distractor’s location wins the competition. This prediction was clearly supported: the compatibility from the difficult distractor was largest when this distractor is cued (M=31%), of intermediate size when the easy distractor was cued (M=19.1%), and smallest when the target was cued (M=14.8%).

Nevertheless, further research with a shorter time interval between the search display and probe onsets may be useful to fully resolve this issue.

### Attentional shifts and attentional engagement

In a recent spatial cueing study, Zivony and Lamy (2018, see also Maxwell, Gaspelin & Ruthruff, 2020) reported both a cue validity effect and an effect of the compatibility between the cued distractor and the target when the cue matched the observer’s attentional set (e.g., a red cue in search for a red target), but only a cue validity effect when the cue did not match this set (e.g., a green cue in search for a red target). The authors concluded that unlike goal-directed capture, stimulus-driven capture elicits “shallow shifts of attention”, that is, attentional shifts (indexed by cue validity effects) that are not followed by attentional engagement (indexed by compatibility effects). The present findings (see also Lamy et al., 2018, Exp.1) are inconsistent with this conclusion: here, the onset cue did not match the attentional set for a circular target, yet this onset cue produced both validity and compatibility effects (in the mixed-difficulty condition).

The Priority Accumulation Framework readily accounts Zivony and Lamy’s findings (see also Lamy et al., 2018, Exp.2^6^) by suggesting that target-matching cues are associated with a larger priority weight than non-matching cues. As a result, when a distractor appeared at the location of a target-matching cue, it was most likely to win the competition (hence the observed cue validity and compatibility effects). In addition, target-distractor similarity was high enough for non-matching cues to bias the competition (hence, the observed spatial cueing effects), but too low for a distractor to win the competition when cued (hence, the null compatibility effects). Although PAF markedly differs from Zivony and Lamy’s (2018) account both in its predictions and in its interpretation of spatial cueing and compatibility effects, it may nevertheless converge with Zivony and Lamy’s (2018) conclusion that goal-directed and stimulus-driven capture of attention have qualitatively different consequences – as explained in the next section.

### What event triggers attentional deployment?

An important, implicit assumption of PAF is that some event must trigger the deployment of attention to the highest-priority location: if attentional priority weights accumulate at each location across time, such accumulation must stop at some point and a signal must indicate when attention should be shifted to the winner. That we should wait for clues indicating that the appropriate moment has arrived for us to deploy our limited attentional resources to the highest-priority location, is a reasonable assumption: it would shield our attention against relentlessly shifting to potentially irrelevant events. The theory does not yet specify what the triggering event might be, yet three main candidates, that are not mutually exclusive, come to mind.

First, temporal expectations may indicate the appropriate time for deploying attention. For instance, in the present study (as well as in Lamy et al., 2018), the temporal sequence was fixed across trials, such that participants could rely on the beginning of the trial, the cue onset, or its offset in order to prepare to deploy their attention in the search display. This suggestion is consistent with previous research showing that temporal expectations are powerful determinants of attentional deployment (see Nobre, 2010 for review).

Second, the onset of the search display itself may signal that the time has come to deploy attention, provided that the search display can be easily discriminated from the events that preceded it. For instance, here, participants may have waited for filled red shapes to appear in order to deploy their attention to the highest-priority location.

A third possibility relies on the findings from previous research showing that feature-based attention is not spatially selective: searching for a feature in a pre-specified region of space improves processing of objects sharing this feature in irrelevant regions of space (e.g., Saenz, Buracas & Boynton, 2002). Such poor selectivity might extend to the temporal domain and detection of the target-defining property might therefore trigger attentional deployment. This hypothesis, if confirmed, would suggest an alternative interpretation of Zivony and Lamy’s (2018) findings. Specifically, in search for a red target, for instance, attention should be allocated to the highest-priority location as soon as a red item is detected. This should occur in the cueing display when the cue shares the target property, but only in the search display when the cue does not share the target property. In the latter case, attention may or may not be allocated to the cued distractor’s location, depending on how similar to the target this distractor is. Thus, this version of PAF preserves the notion that goal-directed and stimulus-driven capture of attention have qualitatively different consequences (in line Zivony & Lamy, 2018) but refutes the claim that attentional engagement occurs only following goal-directed capture.

Future search is required in order to determine what types of events can trigger the deployment of attention.

### Conclusions

The evidence presented here provides novel support for the Priority Accumulation Framework (PAF, Gabbay et al., 2019; Lamy et al., 2018). This model has two central implications for studies of attentional capture. First, it suggests that the question “Does object category X automatically attract our attention?”, which has motivated intense empirical research and theoretical debate, is an ill-posed question. Specifically, the attempt to respond by “yes” or “no” is futile, because whether a given object captures attention depends on the competition context (Desimone & Duncan, 1995). Such context is determined by what other objects surround the critical stimulus, what happened at each location of the visual field in the recent past, what the observers’ goals are, and more. Thus, a spider (e.g., Mogg & Bradley, 2006), one’s own name (e.g., Gronau, Cohen & Ben Shakhar, 2003) or an abruptly onset object (e.g., Yantis & Jonides, 1984) may deserve a special status because they increase the attentional priority accruing to a given location in space relative to other animals, names or luminance changes, but may not reliably trigger a shift of attention to that location.

Second, PAF raises a question that has been largely neglected so far: is attention automatically moved to the ever-changing location with the highest priority, at any given moment? Or does the deployment of attention to the highest-priority location await a trigger that signals the appropriate moment? While most researchers implicitly assume the former to be true, as is clear from the canonical interpretation of spatial cueing effects, PAF assumes the latter. Further research is needed to answer this important issue.

## Author’s note

Support was provided by the Israel Science Foundation (ISF) grants no. 1286/16 to Dominique Lamy.

## Data Availability Statement

The datasets generated during and analyzed during the current study are available in the Open Science Framework (OSF) repository, https://osf.io/3xet7/.

## Footnotes

1. Gaspelin et al. (2016) reported a similar result but did not discuss it.
2. In previous papers (Zivony, Allon, Luria & Lamy, 2018; Zivony & Lamy, 2014; 2016; 2018) we suggested that spatial cueing effects index shifts of attention, whereas compatibility effects index attentional engagement. This alternative view is considered in the General Discussion.
3. We reported an effect of the response compatibility between the difficult distractor and the target when the location of an easy distractor was cued. We initially interpreted this finding as supporting PAF against the Dwelling Hypothesis. However, this finding may simply indicate that after attention was shifted to the cue’s location (where an easy distractor appeared), attention was quickly shifted to the difficult distractor before it was finally redirected to the target, hence the reported compatibility effects between the difficult distractor and the target. This finding is thus equally compatible with PAF and the Dwelling Hypothesis.
4. The search display was not masked. However, as the response feature was a very small black dot, it is highly unlikely that it left a retinal impression, on which participants could have relied to emit a response.
5. In both Gaspelin et al.’s study (2016, Exp.7) and our replications of their experiment (e.g., Exp.1 of the present study), similarity to the target and surface size were confounded, because the target was larger than difficult distractors, which were larger than easy distractors. In addition, when the search display was presented briefly (in Exp.1), some participants reported that in the context of the horizontal ellipses, the perfect circle appeared to be a vertical ellipse. This illusion may have rendered the instruction to look for a perfect circle difficult to follow. To address these issues, all objects were horizontal ellipses occupying the same surface and varying in elongation. Participants were instructed to search for the ellipse that was closest to a circle.
6. There were errors in the report of these results, and these were rectified in an erratum (Lamy, Darnell, Levi & Bublil, 2019).

